# The nanomechanical fingerprint of colorectal cancer -derived peritoneal metastasis

**DOI:** 10.1101/2022.08.17.504271

**Authors:** Ewelina Lorenc, Luca Varinelli, Matteo Chighizola, Silvia Brich, Federica Pisati, Marcello Guaglio, Dario Baratti, Marcello Deraco, Manuela Gariboldi, Alessandro Podesta

## Abstract

Peritoneal metastases (PM) are one of the most common routes of dissemination for colorectal cancer (CRC) and remain a lethal disease with a poor prognosis. The compositional, mechanical and structural properties of the extracellular matrix (ECM) play an important role in cancer development; studying how these properties change during the progression of the disease is crucial to understand CRC-PM development.

The elastic properties of ECMs derived from human samples of normal and neoplastic PM in different pathological conditions were studied by atomic force microscopy (AFM); results were correlated to patients’ clinical data and to the expression of ECM components related to metastatic spread.

Our results show that PM progression is accompanied by stiffening of ECM as a common feature; spatially resolved mechanical analysis highlighted significant spatial heterogeneity of the elastic properties of both normal and neoplastic ECMs, which show significant overlap in the two conditions. On the micrometre scale, ECMs that are considered normal according to the pathological classification possess stiffer spatial domains, which are typically associated with cancer associated fibroblasts (CAF) activity and tumour development in neoplastic matrices; on the other hand, softer regions are found in neoplastic ECMs on the same scales. Our results support the hypothesis that local changes (stiffening) in the normal ECM can create the ground for growth and spread from the tumour of invading metastatic cells.

Mechanical changes correlate well with the presence of CAF and an increase in collagen deposition, which are well known markers of cancer progression. Furthermore, we have found correlations between the mechanical properties of the ECM and patients’ clinical data like age, sex, presence of mutations in *BRAF* and *KRAS* genes and tumour grade.

Overall, our findings suggest that the mechanical phenotyping of the PM-ECM has the potential for predicting tumour development.

## Introduction

Peritoneal Metastasis (PM) affects about one out of every four patients with colorectal cancer (CRC) ^1^. PM development is characterized by several steps where cancer cells disseminate from the primary tumour to the peritoneal cavity ^2^, through a process also known as peritoneal metastatic cascade ^2,3^. To colonize the peritoneum, the neoplastic cells must be able to infiltrate the extracellular matrix (ECM), starting from detachment from the primary tumour, attach to sub-mesothelial connective tissue and receive a favourable host response ^2^.

The ECM is an essential, acellular element of the tissue microenvironment, which plays a crucial role in several processes in tissue homeostasis ^4^. The ECM determines the three-dimensional (3D) structure of the tissue and provides mechanical and biochemical support, playing a major role in cell-cell and cell-matrix communication and cell migration ^4,5^. Moreover, in the last decades, the crucial role of the ECM in cancer progression has been clearly demonstrated. ^6–11^

The extra-cellular microenvironment is composed of water, various fibrous proteins (i.e collagens, elastins, laminins, fibronectins), proteoglycans, glycoproteins, and polysaccharides; the ECM of the specific tissue has unique composition and topology, which results in developing the biochemical and mechanical properties of each organ and tissue ^1,4,5^.

The ECM can be considered a dynamic element of the tissue, as it undergoes several changes in its composition and rearrangements of own components, through covalent and non-covalent modifications, which are associated with cells activity in tissue development, and also severe diseases and cancer progression ^5,12^.

Both mechanical and biochemical changes in the ECM are regulated by growth factors, hormones, cytokines and metalloproteinase (MMP) ^6^. The ECM elastic properties, together with the activity of specific biochemical factors, play a key role in tissue homeostasis, cell fate, cell adhesion, migration, cell cycle progression, differentiation and actin-related cytoskeletal reorganization and contractility ^6,12–14^.

During cancer progression, the ECM undergoes many structural and biochemical changes, such as an increase of collagen deposition, fibres cross-linking and also changes in gene expression^4–6,12–14^. Indeed, the stiffening of the ECM can be observed in pre-malignant and malignant tissues ^6,12^, is associated with high malignancy/aggressiveness and worse prognosis^7,13–15^ and leads to enhanced treatment resistance in most of the tumours ^6^.

Cancer associated fibroblasts (CAFs) can originate from different cell types, including resident fibroblasts and mesothelial cells, which undergo a mesothelial-to-mesenchymal transition (MMT) ^1,16^ and are critical for the progression of the metastatic disease. CAFs oversee the production of ECM proteins such as collagen, fibronectin, and several others as well as proteases and other enzymes involved in post-transcriptional modification of ECM proteins ^16–18^. CAF activation and collagen deposition, which lead to an overall increase of ECM elastic modulus (stiffening) are among the signs of cancer progression. Therefore, the detection of ECM stiffening at the cellular scale could allow us to spot the first signs of tumour development and monitor cancer progression from its beginning. Moreover, a better understanding of the ECM stiffening process and the associated cell-ECM interplay could help develop more efficient therapeutic strategies for the prevention or treatment of cancer.

Atomic force microscopy (AFM) is a powerful and versatile tool to study biological samples at the nano- and microscale, including the quantitative investigation of their morphological and mechanical properties ^19–24^. The mechanical properties of cells, ECMs and tissues can be characterized by AFM ^22–31^ and could constitute a unique mechanical fingerprint of cancer progression ^32^.

Our work started from the hypothesis that the AFM study of nanomechanical properties of cells, ECMs and tissues, when complemented with the analysis of the expression of specific ECM components and with clinical metadata, can provide an important contribution to understanding the mechanisms that lead to the development of the PM. We have therefore studied the changes in the mechanical properties of the peritoneal ECM in patients affected by CRC-PM. In particular, we have characterized the Young’s modulus of elasticity of ECM specimens through indentation measurements performed by AFM ^22^. The results of the nanomechanical analysis have been correlated with CAF presence and collagen organisation in the ECM samples, to obtain information on the physicochemical differences between normal and neoplastic ECMs, and with patient metadata to try to identify mechanical markers related of specific physiopathological state.

## Results

### Changes in the nanomechanical properties of the ECM

The heterogeneity of the ECM samples studied can be appreciated from the violin plots shown in Figure 1A. In several cases, the YM distribution appears as clearly multimodal. It turns out that often the YM value of the highest-order mode is similar (i.e., the distribution shows significant overlap) to the YM value of a leading mode in the distribution of the neoplastic sample (see for example patients 1,2,3,6,7,8).

**Figure 1.**
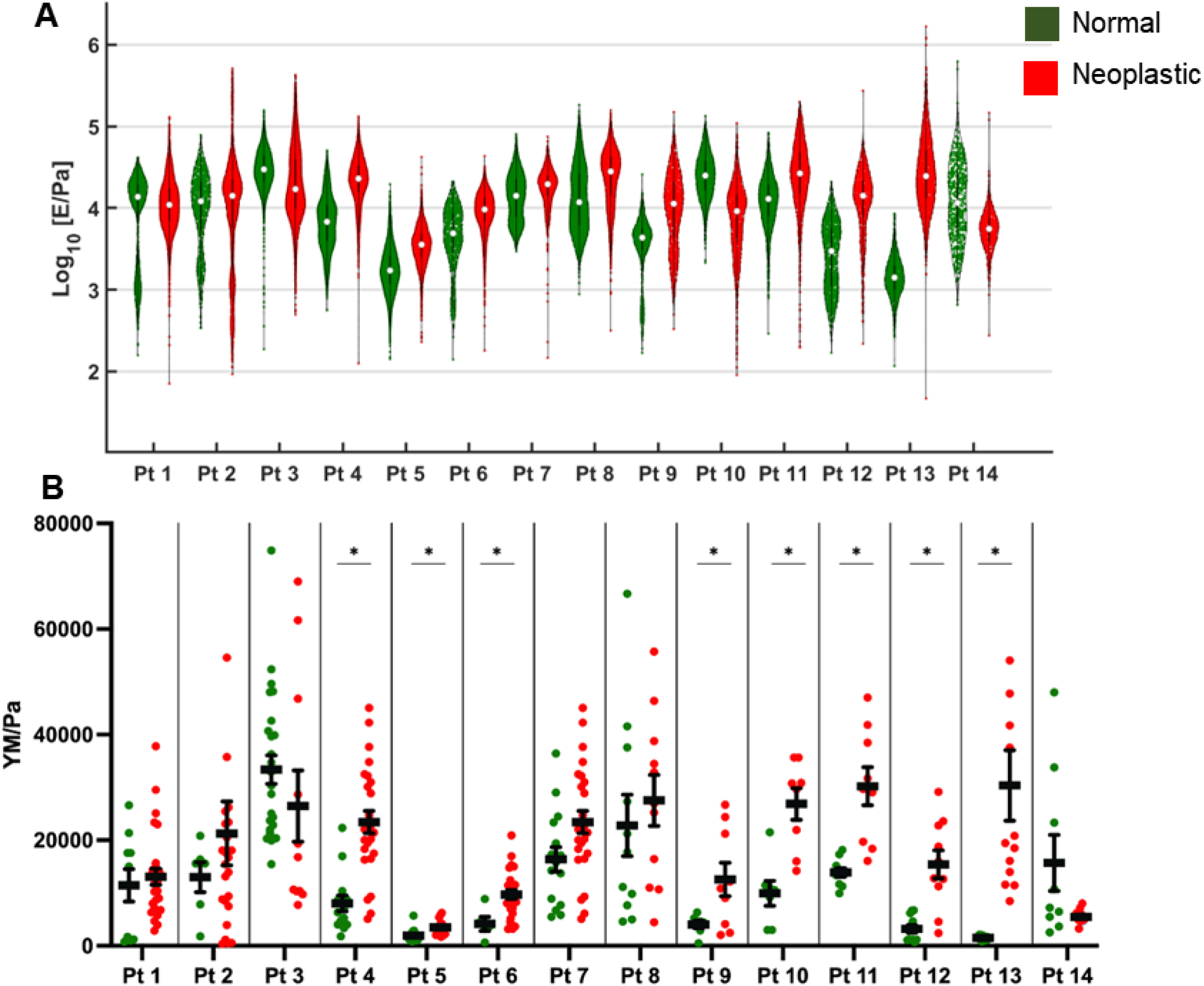
YM distributions for the normal (green) and neoplastic (red) conditions from peritoneal ECMs for the 14 patients considered in the study. A) Violin plots obtained by pooling all YM values from all FCs acquired in all ROIs for a specific condition. The median value is represented by a white dot and black thick lines represent upper and lower quartiles. B) Plots showing the distribution of median YM values measured from all FVs collected in different ROIs for each specific condition. Black dots and bars represent the mean median values and the corresponding standard deviation of the mean, respectively. The asterisk indicates statistical significance of the difference (p<0.05).

Figure 1B shows the distribution of the median YM values measured at different locations of the ECM samples for the tested conditions and patients. In some cases, stiffening is statistically significant, in others it is not, although an increase in the median YM value is often observed. Statistically significant stiffening (i.e., increase of the YM value from the normal to the neoplastic condition) of the CRC-PM derived ECM was observed for eight patients (4-6 and 9-13), who were also among the oldest: 82, 66 and 71 years, respectively, for patients 4-6 and 63, 67, for patients 10-13. However, the stiffening process was also present in patients 7, 8 and 9, who are significantly younger (47, 43 and 45 years old) (Figure 1B).

For both normal and neoplastic tissues, the distribution of YM values is rather broad. Within the same tissue, we observed very wide patient-to-patient variability. For example, for the normal tissue, we observed a difference factor of ∼17 between the YM value of patients 5 and 3; for the neoplastic condition, we observed a difference factor of ∼8 between the YM value of patients 8 and 5. These results highlight, among other aspects, the importance of identifying internal references within the same patient; in our case, this is represented by the normal ECM collected several centimeters away from the cancer lesion.

### Correlation of the mechanical fingerprint with _α_SMA overexpression and collagen fibers presence

To better understand the biological events that sustain the mechanical properties of the ECM, we selected six different cases (patients) and analysed CAF activity related to collagen deposition and orientation by αSMA, and Picrosirius Red staining (Figure 2A,B). Patients whose ECMs exhibit different characteristic mechanical properties (no mechanical differences, moderate and significant stiffening) were selected.

**Figure 2.**
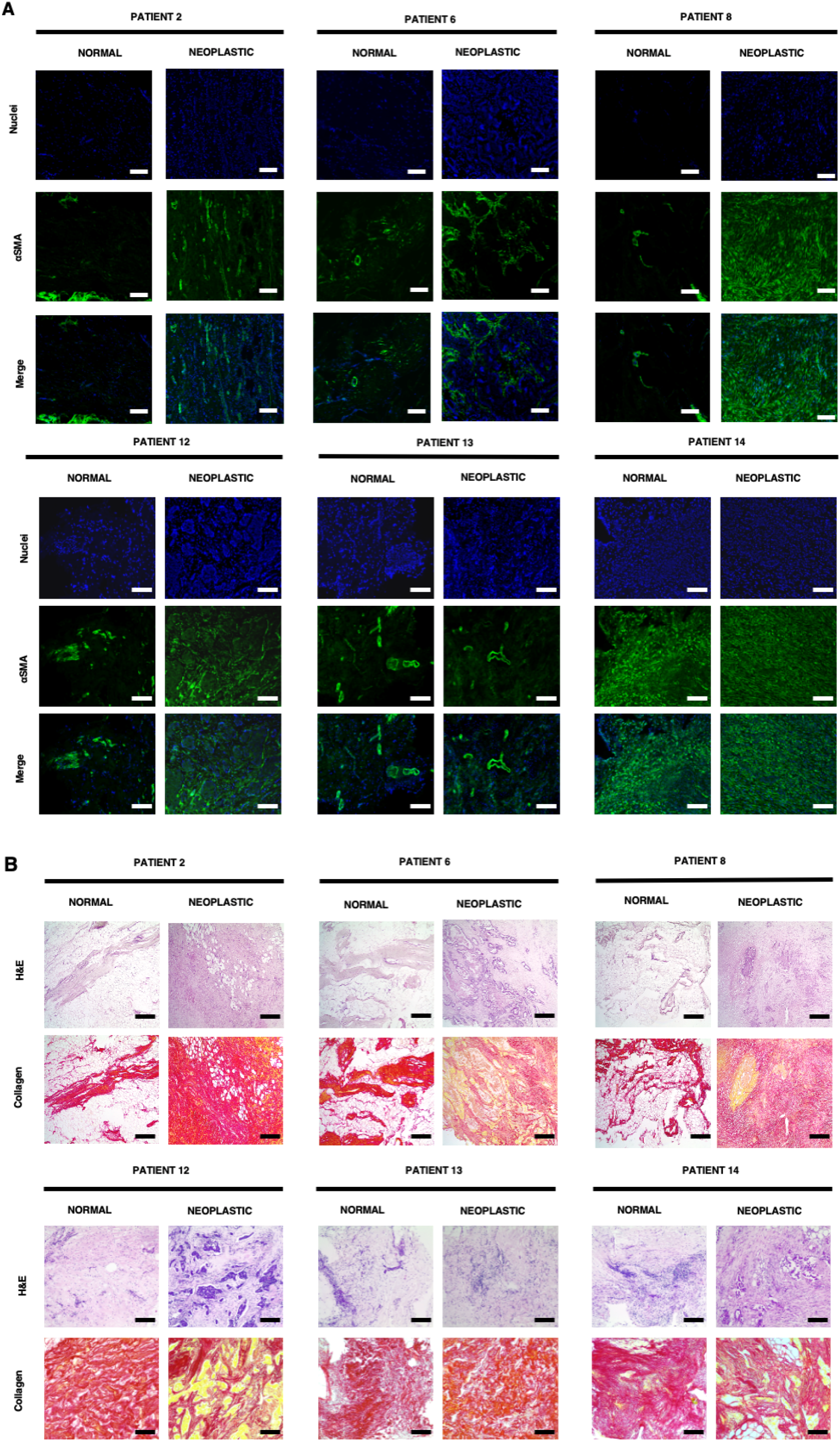
Images of tissue samples from patients 2, 6, 8 and 12-14, for visualization of: A) cell nuclei (DAPI staining) and _α_SMA, magnification 10x; B) collagen and control staining (picrosirius Red and haematoxylin and eosin staining, respectively), magnification 4x.

Five of the six patients showed low expression of αSMA in normal-derived samples, although stromal regions showed high expression of localized αSMA (Figure 2A). Interestingly, we also observed high expression of αSMA around blood vessels (see patients 6, 8 and 13), a sign of the process of neo-angiogenesis, a well-known hallmark of cancer and metastatic spread. Neoplastic-derived samples showed higher expression of αSMA in patients 2, 6, 8, 12; while no clear differences in CAF activity were observed in patients 13 and 14. Patient 13 showed low expression of αSMA in both normal and neoplastic tissues, while patient 14 showed high expression of αSMA in both tissues. For three out of six patients, clear differences in αSMA expression (see Table 1) correlated with ECM stiffening, while patient 14 showed no differences in αSMA and no difference was observed in the measured YM of the two conditions. As can be seen in the case of normal-derived samples, αSMA expression is present, suggesting that neoplastic modifications of the environment are already occurring in a perilesional area (i.e. a region of the tissue that is healthy but close to the tumour mass). In patients 2, 8 and 12 (normal) stiffness distribution was bimodal (Figure 1A) and a similar distribution was observed for αSMA expression.

**Table 1.** Mechano-biological relevant cancer-related modifications. +: Observable differences, ++ marked differences (statistically significant in mechanics), -: no observable differences.

| PATIENTS | αSMA | Collagen | Mechanics | <u>Correlation</u> |
| --- | --- | --- | --- | --- |
| 2 | + | ++ | + | ✓ |
| 6 | + | ++ | ++ | ✓ |
| 8 | ++ | ++ | - |  |
| 12 | ++ | + | ++ | ✓ |
| 13 | - | - | ++ |  |
| 14 | - | - | - | ✓ |

We then evaluated the orientation of collagen fibers by Picrosirius Red staining (Figure 2B). The results showed that normal samples had higher deposition of collagen fibers in stromal and blood vessel areas (Figure 2B), while neoplastic-derived samples were characterized by an irregular and porous orientation of collagen fibers, with a corrugated-like morphology pattern (Figure 2B, see patients 2, 6, 8, 12-14). Overall, collagen results correlated with αSMA expression (see Table 1), demonstrating the active role of CAF in collagen production and deposition during metastatic spread (Figure 2A,B). Again, we observed that some normal-derived samples exhibited a neoplastic-like collagen pattern, also in line with αSMA expression, particularly in patient 14.

### Correlation of the mechanical fingerprint with patients’ clinical data

To better understand how mechanical response and ECM modifications are related to PM, we looked for correlations between the observed biophysical properties and clinical data of the patients.

We first tested whether the Young’s modulus of the normal ECM was correlated with the age of the patients involved, since age-related stiffening has been reported at both the cellular and ECM/tissue level ^33–37^. The results are shown in Figure 3A. It is possible to observe a clear trend toward softening of the normal ECM as the age of patients increases (patient 13, with a colorectal neuroendocrine carcinoma, not included in Figure 3A, showed a decrease in line with the general trend).

**Figure 3.**
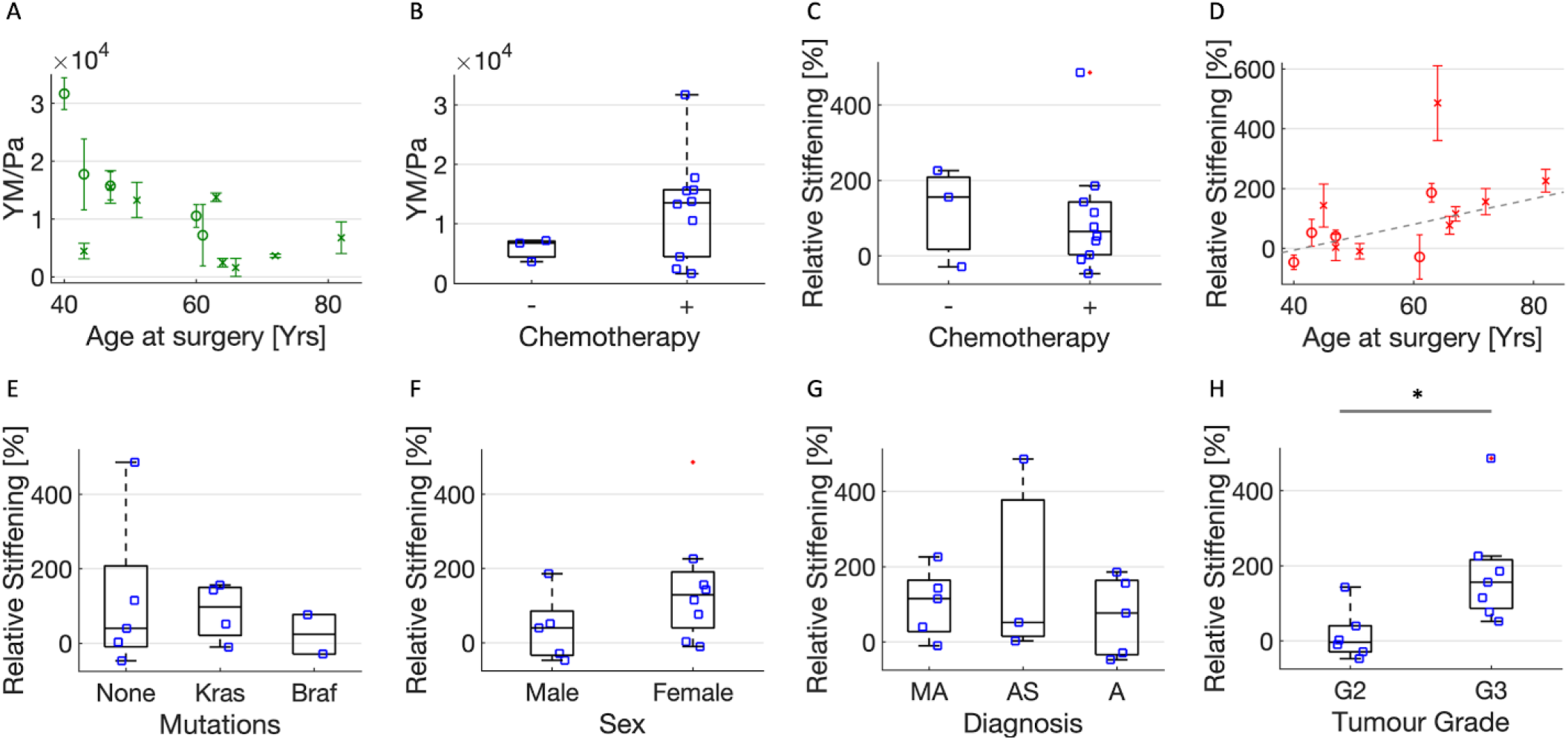
A) YM values (mean median value +/-std of the mean) of healthy ECMs of the 13 patients affected by adenocarcinoma versus their age (circles and crosses represent men and females, respectively). B) YM values as in (A) for patients who did not (-) and did (+) undergo chemotherapy. C-H) Relative stiffening of the neoplastic ECMs from the same patients, versus: C) chemotherapic treatment; D) the patients’ age; E) the presence of mutations in *KRAS* and *BRAF* genes; F) the patients’ sex; G) histology – MA: Mucinous Adenocarcinoma, AS: Adenocarcinoma of the Sigma, A: Adenocarcinoma; H) Tumour grade.

To investigate whether the softening could be due to the treatments undergone by the patients, we checked for correlation with chemotherapy (Figure 3B), but we found no significant evidence of its impact on the elastic properties of the ECM, despite a broader distribution of YM values for patients treated with chemotherapy. To avoid previous treatment-related biases, we calculated the relative stiffening of neoplastic versus normal ECMs within each single patient, as both tissues underwent the same treatments. The relative stiffening was calculated as the difference between the YM values of the neoplastic and normal ECM, normalized to the YM value of the normal ECM. Figure 3C confirms that chemotherapy is not correlated to stiffening of the ECM.

Figure 3D shows that there is a trend toward increasing relative stiffening with the age of the patients, although not significant. Note that patient 13, with a colorectal neuroendocrine carcinoma, showed the strongest increase in stiffness (up to 1200%, and four times larger than the second highest relative stiffening). The very high stiffening observed in patient 13, compared with the other patients, can be a sign of different mechanical modifications between tumour types. However, since neuroendocrine tumours are extremely rare, our observations are not statistically significant and a comparison with the PM group is not possible; patient 13 was therefore excluded from analysis reported in Figure 3.

We tested whether stiffening is related to the presence of mutations in the *KRAS* and *BRAF* genes, which are very frequent in metastatic CRC (Figure 3E). We observed a difference (although not significant) between the relative stiffening of ECMs carrying *KRAS* and *BRAF* mutations, which appeared to be greater in *KRAS* mutated cases. The non-mutated cases spanned a wider range of relative stiffening, encompassing that of the mutated cases.

In addition, we decided to look for correlations between relative stiffening and the sex of patients, since it is known that CRC is more frequent in males than females, although the differences in mortality appear to be insignificant ^38^. Figure 3E shows that females have more pronounced stiffening, although not significant. There are no data on this issue, so it might be interesting to test for sex-related variations in the mechanical properties of other tissues, as in the case of cardiovascular disease, which is known to affect males more than females ^39^.

We also tested whether there are differences in mechanical properties in relation to the histopathological classification of the tumour (Table 2). Figure 3G shows that there are no evident differences in the relative increase of the YM of the ECM of the different histologies.

**Table 2.** Characteristics of the patients from whom the tissue samples were obtained. All samples were at stage IV.

| Patient | Age<sup>1</sup> | Mutations at CRC-related genes<sup>2</sup> | Sex | Diagnosis | Grade | Location | Chemotherapy |
| --- | --- | --- | --- | --- | --- | --- | --- |
| CRC-PM 1<sup>3</sup> | 51 | *KRAS* | Female | Mucinous Adenocarcinoma | G2 | colon | Yes |
| CRC-PM 2 | 47 | NONE | Female | Adenocarcinoma | G2 | sigma | Yes |
| CRC-PM 3 | 40 | NONE | Male | Adenocarcinoma | G2 | colon | Yes |
| CRC-PM 4 | 82 | ND | Female | Mucinous Adenocarcinoma | G3 | right colon | No |
| CRC-PM 5<sup>3</sup> | 66 | *BRAF* | Female | Adenocarcinoma | G3 | right colon | Yes |
| CRC-PM 6 | 71 | *KRAS* | Female | Adenocarcinoma | G3 | colon | No |
| CRC-PM 7 | 47 | *NONE* | Male | Mucinous Adenocarcinoma | G2 | rectum | Yes |
| CRC-PM 8 | 41 | *KRAS* | Male | Mucinous adenocarcinoma | G3 | sigma | Yes |
| CRC-PM 9<sup>3</sup> | 43 | *KRAS* | Female | Mucinous Adenocarcinoma | G2 | right colon | Yes |
| CRC-PM 10<sup>3)</sup> | 60 | ND | Male | Adenocarcinoma | G3 | colon | Yes |
| CRC-PM 11<sup>3)</sup> | 63 | NONE | Female | Mucinous Adenocarcinoma | G3 | colon | Yes |
| CRC-PM 12 | 65 | *NONE* | Female | Adenocarcinoma | G3 | sigma | Yes |
| CRC-PM 13 | 82 | # | Female | Colorectal neuroendocrine carcinoma | # | # | # |
| CRC-PM 14 | 61 | *BRAF* | Male | Adenocarcinoma | G2 | colon | No |

<sup>1</sup> Age at surgery.
<sup>2</sup> *KRAS* and *BRAF* genes were analysed
<sup>3</sup> See Varinelli et al. 2022 <sup>40</sup>.

Finally, we analyzed the differences in the tumour grade of the samples analyzed. Tumour grade describes a tumour in terms of abnormalities of tumour cells compared with normal cells. A low grade indicates a slower growing tumour than a high grade. All patients were diagnosed with grade 2 and grade 3; the results of the correlation between tumour grade and relative stiffening are shown in Figure 3H. It can be observed that the grade 3 group (G3) shows significantly greater stiffening than the grade 2 group (G2).

## Discussion

Changes in the nanomechanical properties of tissues are one of the hallmarks of tumour progression. By understanding the processes behind them, we could exploit this knowledge to implement cancer diagnosis and other diseases based on nano- and microscale mechanical phenotyping. Here, we focused on the investigation of the mechanical modifications in decellularized ECM derived from CRC-PM.

Based on AFM studies on ECM samples from 14 patients, we observed a general trend of ECM stiffening during the development of the neoplastic lesion (Figure 1). Our results agree with previously published data on ECM of other cancer types ^10,40,41^.

The measured multimodal, not simply broad, distributions of YM values, suggest that the ECM possesses significant structural, compositional, and therefore also mechanical heterogeneity at the AFM measurement scale (10-100 μm). Moreover, the partial overlap of YM values distributions of different conditions (normal and neoplastic), revealed the complexity of disease progression during the metastatic process, which is characterized by high spatial heterogeneity at the cellular and supracellular scale. The seed and soil theory suggests that ECM undergoes changes, including mechanical ones, to prepare a microenvironment suitable for neoplastic cell proliferation ^42^. The presence of stiffer regions in normal samples, comparable to those typical of the neoplastic cases, suggests that local changes that prepare the ground for metastatic invasion in the normal tissue occur far from the existing lesion, likely caused by the release of factors that can ultimately alter the mechanical properties of the ECM ^42^. The common practice for cancer studies is to obtain non-tumoural sample 10-15cm away from the tumour ^26^. Our results on the mechanical properties of the ECM show that the practice of considering the tissue located 10-15 cm far from the cancer lesion as non-tumoural can be uncorrect.

Staining for αSMA, a CAF-specific protein expressed in fibroblasts, which is a sign of cancer progression and a typical marker of desmoplasia ^16^, detected changes in the surrounding microenvironment that typically lead to the development of specific metastatic niches. Since we also observed areas with high expression of αSMA in normal samples, it is likely that these may have undergone modification induced by specific pro-metastatic factors released by nearby PM metastatic cells. These results confirmed that the formation of pre-metastatic niches already occurs in normal derived tissue as we also observed areas of high αSMA expression in normal samples.

Higher expression of αSMA was also observed in areas rich in blood vessels (patients 6, 8 and 13). During the metastatic process, CAFs can distribute through blood vessels to develop the so-called perivascular metastatic niches, which can sustain the activation of stromal cells in normal tissue through TGF-ß and secretion of pro-inflammatory and pleiotropic interleukins and cytokines, which also contribute to initiate modifications in the ECM, including stiffening. The combination of these events generates a microenvironment more suitable to support metastatic spread, in particular angiogenesis ^16,18,42^. Expression of αSMA in vascularized areas indicates early steps of metastatic invasion into normal tissues.

Expression of αSMA was higher than normal tissue in four of the six patients and CAF activity correlated with increased collagen deposition; nevertheless, two patients (8 and 13) exhibited uncorrelated results between staining and mechanical differences of normal and neoplastic ECM. Based on these results it appears that some of the tissue that was considered normal already had cancerous characteristics.

Another step to better understand the differences between normal and neoplastic ECM was to visualize collagen I and III fibers, as their overexpression and remodelling are strictly related to cancer progression^4–6,23,43^. Our results showed the typical expression and organization of collagen fibers already observed in previous studies on different types of ECM^10,30,44^. The organization of collagen fibers was strictly correlated with the expression of αSMA; expression of this protein confirmed collagen crosslinking in neoplastic ECM but also to a smaller extent in normal ECM. Tumours with high desmoplasia (fiber crosslinking) are considered more aggressive and with a worse prognosis ^6^. In the neoplastic samples, increased crosslinking and restructuring of collagen fibrils in the ECM, and matrix stiffening produce an extracellular environment conducive to tumour invasion and growth. Changes in the vascularized and stiffer perilesional area could be a feed-forward loop to spread neoplastic ECM characteristics. Myofibroblasts are known for their ECM remodelling, which involves *de novo* deposition of specific receptors involved in mechanosignaling by the ECM, contributing to both normal and pathologic tissue remodelling ^45^.

Correlation of these results with patients’ clinical data suggests a clear trend for tissue softening in older patients. This result is somewhat unexpected, considered that age-related stiffening at both cells and ECM/tissue levels has been reported ^33–37^; damaging and inflammation of the tissue due to an extended inflammatory condition related to the presence of the tumour ^28^ could explain our observation. We exclude that the softening can be directly related to chemotherapy treatments, since we did not find any evident correlations in our patients (Figure 3B,C).

We then focused the analysis on the relative stiffening of neoplastic versus normal ECMs in each individual patient. The analysis of the mechanical properties according to the mutational status of *KRAS* and *BRAF* genes and across tumour grade showed that patients with mutations in *KRAS* gene had a slightly higher relative stiffnening, while a stronger relative increase is related to different tumour grade (G3 > G2). Since the presence of mutations in *KRAS* and *BRAF* genes is very common in PM and tumour grade is a parameter that characterizes tumour cell behaviour, they are probably associated to specific mechanical characteristics. These data are still preliminary and will be investigated with further experiments on a larger cohort of cases. We believe that such correlations would help to advance the development of biomechanical tests to complement standard clinical diagnostic techniques.

Understanding how modifications of the mechanical properties of the ECM influence the metastatic invasion may also have the potential in developing active tissue treatments that can impact on cell migration; ECM is already being used as a scaffold for cell culture to better understand the cell-microenvironment interaction mechanisms ^23,40,46–50^.

In conclusion, we provided evidence that in CRC-PM ECM stiffening correlates with collagen deposition and remodelling, CAF activity, age at surgery and tumour grade. Spatially resolved mechanical analysis of human-derived samples revealed significant spatial heterogeneity in the elastic properties of normal and neoplastic ECMs. The results, together with the high expression of αSMA, revealed that signs of pre-metastatic niche formation are already present in normal tissue, and correlation of mechanical data with patients’ metadata showed interesting connections between the relative stiffening and characteristics of the tumour itself, in particular with patients’ age and tumour grade. Our results suggest that nano- and microscale characterization of tissue mechanical properties can suggest the presence of metastasis and help in diagnostic procedures. A better comprehension of the mechanical properties of ECM will facilitate its use as a scaffold for culturing cells in future research, creating more reliable models of the disease.

## Materials and methods

### Sample preparation

ECMs were obtained from the peritoneal tissue of 14 patients diagnosed with CRC-PM (more detailed information is in Table 2).

The samples were collected during surgical resection at the Peritoneal Malignaces Unit of Fondazione IRCCS Istituto Nazionale Tumori di Milano as described in Varinelli *et al*. ^40^ The study was approved by the Institutional Review Board of Fondazione IRCCS Istituto Nazionale Tumori di Milano (134/13; I249/19) and was carried out following the Declaration of Helsinki, 2009. All experiments were performed in accordance with relevant named guidelines and regulations. Written informed consent was obtained from all participants.

Briefly, non-tumoural tissues were collected 10 cm away from tumour, according to standard clinical procedures.^26^ Neoplastic-derived and normal-derived 3D decellularized extracellular matrix (3D-dECM) specimens were obtained as described in Genovese et al. ^51^. 3D-dECM were embedded in OCT and then frozen in a liquid nitrogen bath of isopropanol. Frozen samples were cut into slices of 100-200μm thickness and immobilized on polarized glass slides (Thermofisher, Walthan, USA). Prepared samples were stored at -4°C and used for the AFM measurements.

### AFM nanoindentation measurements

The nanomechanical measurements were performed using a Bioscope Catalyst AFM (Bruker) mounted on top of an inverted microscope optical microscope (Olympus X71). To isolate the AFM from ground and acoustic noise, the microscope was placed on an active antivibration base (DVIA-T45, Daeil Systems) inside an acoustic enclosure (Schaefer, Italy). AFM measurements of cryosections were performed in a PBS droplet confined by a circle of hydrophobic ink.

AFM-based nanomechanical measurements were performed according to standard procedures, based on the acquisition of indentation curves, as described in Refs ^22,52^. The measurement process is described schematically in Figure 4. We have used custom colloidal probes with spherical tips made of borosilicate glass beads with diameter (twice the radius R) in the range 18-25μm, produced and calibrated as described in Ref. ^53^. The spring constant of the AFM probes (typically 5-6 N/m) was calibrated using the thermal noise method ^54,55^. The deflection sensitivity of the optical beam deflection apparatus (in units of nm/V) was calculated as the inverse of the slope of the force vs. distance curves (simply force curves, FCs) collected on a stiff substrate (the glass slide holding the sample)^52^ or using the contactless SNAP procedure^56^.

**Figure 4.**
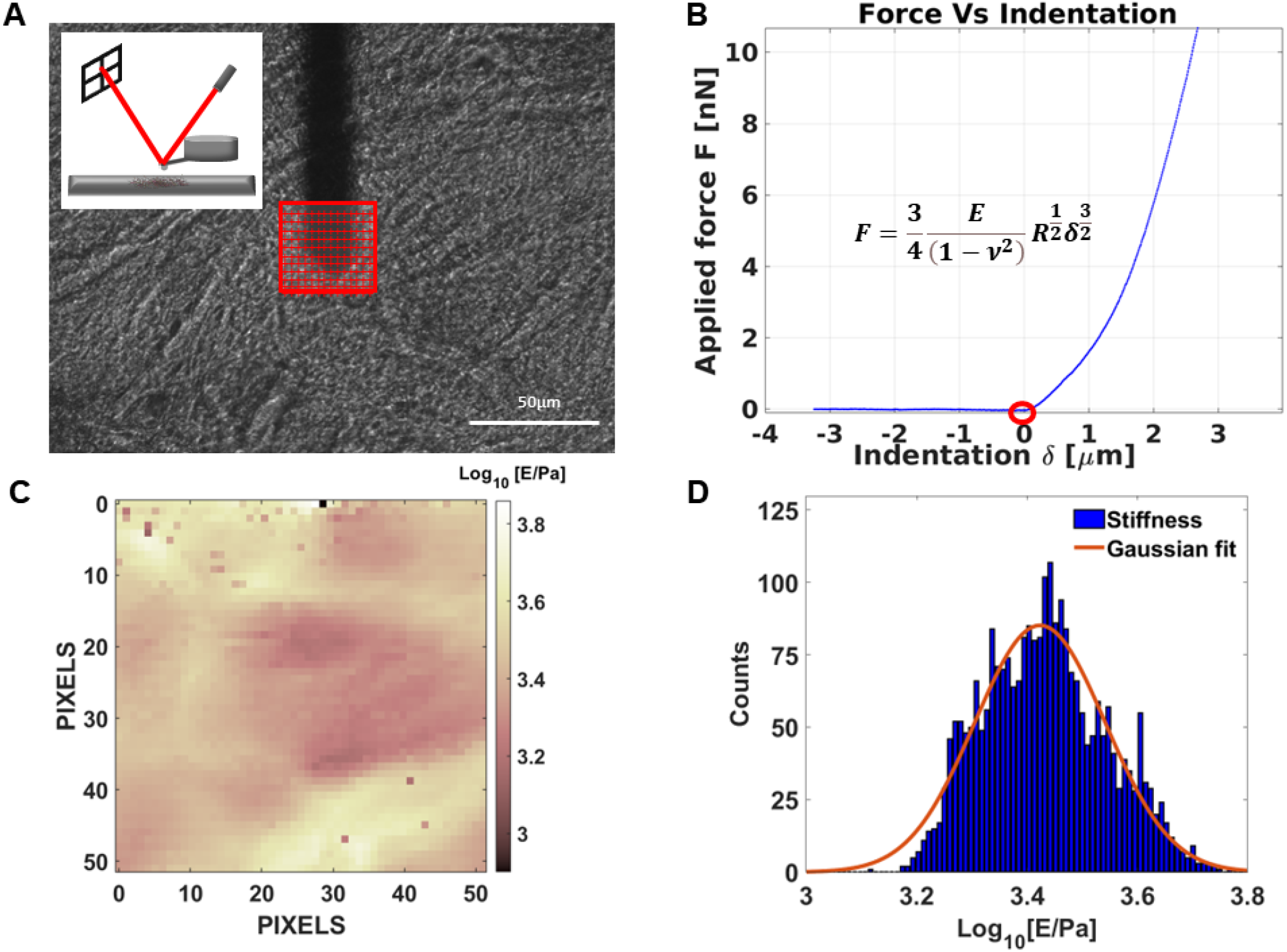
Schematic representation of the nanomechanical measurement. A) Optical image of an ECM (normal peritoneum-derived from patient 8), with the AFM cantilever and the selected region of interest for the indentation experiment. Slices with thickness between 100 and 200 μm are semi-transparent, which allows to select regions for the analysis that look sufficiently uniform and smooth. The red grid represents the locations where force curves (FCs) are acquired (scale bar length – 50μm). In the inset, the experimental setup for indentation measurements and the optical beam deflection system are shown. B) Typical rescaled approaching force vs indentation curve. The red circle highlights the contact point. Only the portion of the curve characterized by positive indentation is considered for the Hertzian fit (Eq. 1 and also shown in the inset). C) The map of Young’s modulus values extracted by the FCs acquired in the region of interested shown in A. D) Histogram representing the distribution of YM values represented in the mechanical map in C. Under the hypothesis of a log-normal distribution, a Gaussian fit in semi-log scale allows to identify the median YM value, as the centre of the Gaussian curve.

The samples were studied by collecting set of FCs, also called force volumes (FVs), in different regions of interest (ROIs). Each FV typically covered an area between 50μm x 50μm to 125μm x125 μm and consisted of 100-225 FCs. The separation between adjacent FCs was chosen to be greater than the typical contact radius at maximum indentation, to reduce correlations between neighbouring FCs. For each patient’s condition (normal or neoplastic), several FVs were collected on different, macroscopically separated locations on each sample, with 2-3 samples (cryosections) per condition (normal vs neoplastic) for each patient. In total, for each condition 2000-4000 FCs were collected. A FC typically contained 8192 points, with ramp length L = 15 μm, maximum load F_max_ = 800 – 1500 nN, ramp frequency f = 1 Hz. The maximum load was adjusted to obtain a maximum indentation of 4-6 μm in all samples.

Acquired data was analysed in MATLAB using protocol previously described in Puricelli et al.^22^. The elastic properties of the ECMs were characterized through their Young’s modulus (YM) of elasticity, extracted by fitting the Hertz model ^57,58^ to the 20%-80% indentation range of the FCs (details in Ref. ^20,22^):

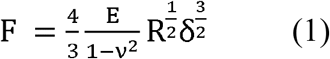

, which is accurate as long as the indentation δ is small compared to the radius R. In Eq. (1), ν is the Poisson’s coefficient, which is typically assumed to be equal to 0.5 for incompressible materials, and E is the YM.

Finite thickness correction ^22,59–61^ was not applied since the thickness of the ECM slices (150-200μm) is significantly larger than the expected contact radius at maximum indentation. The first 20% of the FCs is typically ignored, due to the contribution of superficial non-crosslinked fibers, surface roughness issues, etc.^10^.

### Histochemistry (HC) and immunofluorescence (IF) analyses

Before HC and IF staining, formalin-fixed-paraffin-embedded (FFPE) blocks were prepared and cut as in Varinelli *et al* ^40^. FFPE sections were stained with haematoxylin and eosin (H&E) for visualized nuclei and stromal regions. For HC analysis, sections were stained with picrosirius red (ScyTek lab), to visualize collagen fibers, following the manufacturers’ instructions. Antigen retrieval IF analysis was carried out as in Varinelli *et al*^40^. For IF analyses, FFPE sections were stained with primary alpha Smooth Muscle Actin (αSMA,1:400), FITC conjugated antibody (Merck, KGaA) and DAPI (Merck, KGaA), following the manufacturers’ instructions, to visualize respectively cancer associated fibroblasts (CAF) and nuclei. Images were acquired with a DM6000B microscope (Wetzlar, Germany Leica,) equipped with a 100 W mercury lamp, and analyzed using Cytovision software (Leica). All the experiments were performed in triplicate.

### DNA sequencing

Presence of mutations in *KRAS* and *BRAF* genes was determined by DNA sequencing. DNA from FFPE sections of PM patient’s tissues was used for mutational analysis and extracted as in Varinelli *et al*^40^. About 150–200 ng genomic DNA (measured with Qubit dsDNA HS assay kit, ThermoFisher Scientific), were sheared by the Sure Select Enzymatic Fragmentation kit (Agilent Technologies Inc., Santa Clara, CA, USA). NGS libraries, probe set design, DNA sequencing and data analysis were performed as in Varinelli *et al* ^40^.

### Statistics

To evaluate the distribution of YM values that is peculiar of ECM in the different tested conditions, all values of the YM from all FCs were pooled together. This is justified in part by the fact that curve to curve distance is of the order of the maximum contact radius, and by the fact that separation between FVs from the same slice is comparable to the separation between FV from different slices obtained from the same patient. To highlight the diversity of local mechanical conditions met in the samples, we have used violin plots to represent the YM distributions (Figure 1A).

The evaluation of representative YM values for a specific condition of a specific patient has been done by pooling the median YM values obtained from all FVs collected in different ROIs and calculating their mean value and the corresponding standard deviation of the mean^28^, assuming that the median values should be normally distributed according to the central limit theorem ^62^. The distributions of median values are shown in Figure 1B. An experimental relative error of approximately 3%, evaluated using a Monte Carlo method ^63^, taking into account the uncertainties in the calibration factors (10% for the spring constant, 5% for the deflection sensitivity) was added in quadrature to the standard deviation of the mean to estimate the final error associated with the mean median YM values.

The statistical significance of differences between tested conditions was assessed using a two-tailed t-test. In case of a p-value <0.05, the difference was considered as statistically significant.

## Data availability

The datasets generated and/or analysed during the current study are available from the corresponding authors on reasonable request.

## Acknowledgements

This research was funded by the European Union Horizon 2020 research and innovation program under the Marie Skłodowska-Curie grant agreement No. 812772, project Phys2Biomed, and under FET Open grant agreement No. 801126, project EDIT. This work was supported by “5 per 1000” funds (2019 MUR and 2015 Ministry of Health, financial support for research) – institutional grant BRI 2021 “Harnessing the extracellular matrix to awaken the immune response in patients with peritoneal metastasis” assigned to Dr. Luca Varinelli, and by the Italian Ministry of Health, with a grant agreement No. RF2019-12370456. We thank the patients who participated in the study. We thank Hatice Holuigue for her valuable support and discussions.

